# *Acinetobacter* spp. with lower susceptibility to quaternary ammonium compounds enriched in microbial communities of frequently used sinks

**DOI:** 10.1101/2025.09.29.679312

**Authors:** MC Cruz, R Ruhal, J Lavin, S Bridwell, N Maghboli Balasjin, B Raasch, R Melton, BK Mayer, CW Marshall, K Hristova

**Affiliations:** Department of Biological Sciences, Marquette University, Milwaukee, WI, USA; Instituto de Investigaciones para la Industria Química (INIQUI), Universidad Nacional de Salta (UNSa) – Consejo Nacional de Investigaciones Científicas y Técnicas (CONICET), Salta, Argentina; Department of Civil, Construction and Environmental Engineering, Marquette University, Milwaukee, WI, USA

## Abstract

Sanitary environments that undergo frequent cleaning and disinfection may harbor microbial communities with potential health risks. While biofilms in healthcare settings are well studied, comparatively less is known about sink microbiomes in public and educational buildings, where hundreds of people may interact with shared sink fixtures. This study characterized the spatial and temporal heterogeneity of sink drain biofilm microbiomes in academic buildings. We sampled 16 sinks from two buildings (four floors each, with sinks closest and furthest to the bathroom entrance), which are cleaned daily with quaternary ammonium compound (QAC) disinfectants, during periods of low and high student traffic (during and after academic breaks, respectively) across winter, spring, and summer. We observed significant spatial and temporal variations in microbial assemblages. Individual sinks accounted for 43% (PERMANOVA, *p < 0.0001*) of the variation in microbial communities. Microbiomes in each building were dominated by two genera, which together accounted for 30% of the community composition: *Acinetobacter* and *Enhydrobacter* (also classified as *Moraxella*) in the newer building, and *Sphingomonas* and *Mycobacterium* in the older building. *Acinetobacter* abundance varied seasonally and showed higher relative abundance during periods of high traffic. Metagenomic analysis of selected sinks revealed a high prevalence of *qac* genes and metagenome-assembled genomes (MAGs) harboring antimicrobial resistance genes (ARGs) including *A. parvus.* Notably, 34–53% of *qac* genes were co-localized on contigs associated with mobile genetic elements. These findings suggest that disinfected sink drains serve as persistent reservoirs of diverse microorganisms and potentially mobile resistance elements.

**IMPORTANCE:** Sink drains are recognized environmental reservoirs for multidrug-resistant bacteria and have been linked to healthcare-associated outbreaks. In public and educational buildings, these microbiomes are shaped by frequent human activity, making them potential sources of exposure and contributors to the environmental dissemination of antibiotic resistance genes. Quaternary ammonium compound (QAC) disinfectants are widely used on surfaces; however, they can select for resistant taxa and co-select for antibiotic resistance. In this study, despite routine cleaning with QACs, public restroom drains remain colonized by resilient biofilms, posing a potential risk to multiple users. Additionally, factors such as human traffic and seasonal variation may influence drain usage and microbial community composition. Elucidating how seasonal dynamics and human activity shape sink biofilms is essential for understanding their role in the environmental transmission of antimicrobial resistance and informing mitigation strategies in nonclinical settings.

## INTRODUCTION

Sink drains have been identified as important reservoirs for multi-drug-resistant bacteria (Decker & Palmore, 2013) and can be a source of infection linked to outbreaks and environmental transmission in the healthcare environment (Brooks et al., 2017). However, there is limited research on non-healthcare settings like residences and public academic buildings (public restrooms, dormitories), where thousands of people spend several hours daily (Flores et al., 2011; Gibbons et al., 2015; Lax et al., 2017; Miles et al., 2019; Qureshi et al., 2020; Richardson et al., 2019; Ross & Neufeld, 2015; Sharma et al., 2019). In high-income urban areas, where populations typically spend approximately 90% of their time indoors (Klepeis et al., 2001), the built environment and its associated indoor microbiome serve as a significant niche for environmental microorganisms (Gilbert & Hartmann, 2024). These microbial reservoirs play a crucial role in fostering and shaping the human microbiome, even from an early age (Brooks et al., 2018; Brooks et al., 2014). Buildings host complex microbial communities that play a pivotal role in shaping and maintaining a healthy human microbiome and are directly and interdependently influenced by social, spatial, and technological factors (Browne et al., 2017). This relationship is becoming increasingly critical, as it is estimated that by 2050, two-thirds of the global population will reside in urbanized built environments (United Nations Department of Economic and Social, 2019). Thus, the design of modern buildings should consider the influence of microbiomes, and research on host–microbiome interactions should increasingly incorporate the role of the built environment as a key factor (Bosch et al., 2024).

Indoor environments, which are constantly exposed to cleaning and disinfection protocols, serve as reservoirs of environmental microbes (Pausan et al., 2022). Cleaning and disinfection protocols can lead to microbial adaptations that pose health risks, such as the development of antibiotic resistance, highlighting the need for careful management of microbial ecosystems within the built environment (Hu et al., 2023). Moreover, the widespread use of disinfectants, particularly quaternary ammonium compounds (QACs) like benzalkonium chloride (BAC) commonly present in household cleaners, has raised concerns about their potential contribution to the development and increase of antibiotic resistance (Mohapatra et al., 2023). Surface attached microbial communities and populations in biofilms exhibit significantly greater resistance to biocides compared to planktonic cells, allowing bacteria to persist on surfaces even after disinfection (Maillard & Centeleghe, 2023). Therefore, it is important to investigate the effects of QAC disinfectant exposure on the dynamics and antimicrobial resistance of biofilm communities in public premise plumbing systems and bathrooms.

Sink drains provide an ideal habitat for the formation of microbial biofilms, as they are consistently moist, often humid, and well-protected environments (Butler et al., 2022). The steady influx of nutrients, introduction of bacteria from users’ hands, disposal of fluids, and daily cleaning and disinfection efforts all contribute to the development of microbial communities that can host opportunistic pathogens, including disinfectant and antibiotic-resistant bacteria (Franco et al., 2020). Moreover, opportunistic pathogens or mobile genetic elements (MGEs) harboring antibiotic resistance genes (ARGs) lead to increased exposure risk (Martínez et al., 2015). Likewise, the spread and propagation of microbes from sink drains via airborne aerosols can contribute to indoor transmission of diseases (Döring et al., 1996).

Microbial heterogeneity is primarily driven by both spatial and temporal differences in environmental factors, and it is reasonable to anticipate spatial heterogeneity within the building premise plumbing system (Neu et al., 2019). These differences are not confined to distinct sections of a system or different building floors but can also manifest within individual fixtures or adjacent sink drains (Withey et al., 2021). It is likely that the same conditions affecting the growth of heterotrophic bacteria also impact the growth of pathogens. A previous study conducted in the UK analyzed 123 samples from two parts of sinks (P-trap and below-strainer) across nine buildings on a university campus (Withey et al., 2021). The study found that microbial community structure and compositions were highly variable across individual sinks, with marginal differences between the two sections sampled and buildings. We wanted to explore this further and determine how seasonality and variation in foot traffic relative to building occupancy levels would change the sink drain microbial communities.

The goal of this study was to characterize the spatial and temporal heterogeneity of biofilm microbiomes in sink drains across academic university buildings. We hypothesized that i) sink drains have distinct microbial communities among locations and through time, and ii) microbial communities in sink drains would harbor genes associated with disinfectant resistance due to frequent use and cleaning. To test these hypotheses, we conducted a study in two academic buildings, systematically sampling two sinks per restroom -one located closest and the other furthest from the entrance- on each of four floors, resulting in a total of eight sinks per building. We monitored sink drain microbiomes during periods of low and high student traffic (during and after academic breaks, respectively) over three seasons (winter, spring, and summer). To test the second hypothesis, we used metagenomics to evaluate the presence of *qac* genes, ARGs, and their relationships with MGEs within biofilm samples coupled with cultured communities grown on media plates containing varying concentrations of disinfectants.

## 2. MATERIAL AND METHODS

### 2.1 Sample collection

We selected two academic buildings on the Marquette University campus in Milwaukee, WI, WC and EH, constructed in 1960 and 2011, respectively. We monitored 16 sink drains, eight in each building. In both buildings, sinks were located in women’s restrooms across four levels, with two sinks selected in each restroom: one positioned closest to the entrance (A) and the other furthest from the entrance (B). In the newer EH building, each restroom had four sinks, with the soap dispersers centrally located between sinks and the paper towel dispensers near the door. In the older WC building, the number of sinks varied by floor. The basement had three sinks, the first level had four sinks, and the second and third levels each had two sinks. In these restrooms, the soap disperser was equidistant from the sinks.

Samples were taken over a nine-month period from January to September 2022, encompassing three seasons: winter, spring, and summer (Table S1). In each season, two sampling campaigns were conducted, one during low traffic and one in high traffic. Periods of low and high traffic were determined based on the academic calendar, with sampling scheduled during class breaks (low traffic) and active semester weeks (high traffic). During the summer season, traffic was quantified using an infrared beam sensor placed at the entrance of the restroom of each floor for four consecutive days, during the summer break and two weeks after classes had resumed (Table S2). These people-counting sensors became available only toward the end of the sampling during the summer campaign and were used to compare the expected difference in traffic between break and class periods. In the WC building, differences were observed only on the first floor where large auditorium classes are located and teaching labs are placed, the other floors are used mostly by professors and occupants of research laboratories. In the EH building, differences were observed in the first, second, and fourth floors. In this building, seminar rooms, teaching and research labs are distributed on every floor.

Replicate swab samples were collected three times per day. Two in the morning, and another sample in the afternoon, after 8 h of restroom usage during class time. These three samples were used as replicate samples for each sink in our subsequent analyses. In total, 288 swab (ESwab collection and transportation system, BD) samples were collected from sink biofilms. Daily cleaning of the sinks in both buildings was conducted throughout the entire sampling period, except on weekends. The cleaning process involved spraying a disinfectant cleaner (Virex^®^Plus, diluted 1:256) and wiping down the tap and sink surfaces. This was performed late at night in EH (10 pm) and early in the morning in WC (6 am). Furthermore, the soap used in the dispersers contained a green-certified foam skin cleanser composed of surfactants and foaming agents (sodium laureth sulfate, cocamidopropyl betaine, PEG-7 glyceryl cocoate), humectants agents (propylene glycol, glycerin, panthenol), chelating and buffer agents (sodium citrate, citric acid), and preservatives with antimicrobial properties (methylchloroisothiazolinone and methylsothiazoline).

Many studies suggest that the microbiota of water-related indoor habitats, like sinks, primarily originates from tap water (Novak Babič et al., 2020). To analyze whether incoming bulk water continuously inoculated downstream sink drains, we obtained composite water samples from sink taps at each floor of the academic buildings in parallel to biofilm samples. A volume of 12.5 L of water was sampled in both the morning (AM) and in the evening (PM). In total, we collected 24 composite samples (12 per building). Bulk water samples were analyzed for physiochemical and chemical parameters. Water quality parameters measured included pH, temperature, and metal ions quantified by inductively coupled plasma mass spectrometry (Agilent 7700x ICP-MS with ASX-500 autosampler). Prior to quantification, samples were filtered through a 0.45 μm membrane and acidified with 2% HNO_3_ and 0.5% HCl (v/v) and stored at 4 °C until analysis according to Method 3125B (APHA, 2017).

### 2.2 Exposure to BAC and Virex disinfectants

To assess the ability of sink drain bacteria to grow on BAC-containing media, we performed cultivation assays using suspensions prepared from swab samples collected from the two academic buildings (EH and WC). These suspensions were diluted and spread onto R2A plates containing varying concentrations of BAC (C_14_ 50%, C_12_ 40%, C_16_ 10%) at 1, 10, and 100 mg/L, as well as to Virex^®^Plus (Diversey, Inc., Charlotte, NC), the commercial disinfectant routinely used for sink cleaning. Virex^®^Plus was tested at two dilutions: V1 (1:16,000 times), corresponding to a BAC concentration of 5.4 mg/L, and V3 (1:2,000 times - 8 times more diluted than recommended), corresponding to 43.5 mg/L BAC. In addition to BAC, the commercial disinfectant formulation included other QAC biocides: 6.67% octyl decyl dimethyl ammonium chloride, 2.67% dioctyl dimethyl ammonium chloride, and 4% dodecyl dimethyl ammonium chloride. Plates were incubated at room temperature for seven days. After incubation, all colonies from the plates were collected using a sterile loop, and the resulting pellet was preserved at -20 °C until DNA extraction.

### 2.3 Bacterial DNA extraction

Bacterial DNA was extracted directly from swab tips plus 250 μL of residual collection buffer (BD ESwab collection and transportation system) using the QIAamp PowerFecal Pro DNA kit (Qiagen, Germany) following the manufacturer’s protocol. We used a homogenizer (Bertin Technologies, France) for sample bead beating for two cycles of 60 sec and 10 sec pause as a first step before proceeding with the Qiagen protocol. Extracted DNA was dissolved in 80 μL sterile nuclease free water and stored at −20 °C until further analyses. Extraction blanks were included for each new kit used.

Each water sample (12.5 L) was filtered using a 0.22 μm polycarbonate filter (Millipore, Germany). Subsequently, all filters were stored at -20 °C until DNA extraction. Extraction of the DNA from water samples was performed as described by Beattie et al. (2018). Briefly, filters were added to 250 μL lysis buffer followed by three cycles of freeze and thaw with liquid nitrogen. Later, 250 μL was added to zirconium beads (0.1 mm), followed by cell disruption by bead beating for 60 sec at 2100 oscillations/min using a Mini-Bead Beater (BioSpec Products, Bartlesville, OK, USA).

Extracted DNA samples were sent to the Environmental Sample Preparation and Sequencing Facility at Argonne National Laboratory for 16S rRNA amplicon sequencing. Reagent and extraction blanks were included for each batch of DNA extraction to ensure no carryover contamination occurred during DNA extraction. No carryover contamination was observed in reagent blank samples. Furthermore, genomic DNA from swab and culture plate samples was extracted for metagenomic analysis using whole genome sequencing.

### 2.4 16S rRNA amplicon and whole genome sequencing

Amplicon sequencing was performed using primers designed to be massively multiplexed and cover the V4 hypervariable region of the 16S rRNA gene (515F-806R) using the standard methods outlined by the Earth Microbiome Project (EMP) (http://www.earthmicrobiome.org/emp-standard-protocols/16S/). Samples were sequenced on the Illumina MiSeq platform, at the Environmental Sample Preparation and Sequencing Facility (ESPSF) at Argonne National Laboratory.

Samples were prepared for whole metagenome sequencing using an Illumina DNA PCR-free Prep tagmentation kit and unique dual indexes. Sequencing was performed on the Illumina NextSeq2000 platform using a 600-cycle flow cell kit to produce 2x300 bp paired reads. A 2% PhiX control was spiked into the run to support optimal base calling. Metagenome samples were sequenced at SeqCoast Genomics.

### 2.5 Bioinformatic analysis

*16S rRNA data analysis* We performed quality filtering, merging of paired reads, and amplicon sequence variant (ASV) clustering using DADA2 package version 1.28.0 (Callahan et al., 2016) in R version 4.3.3 (R Core Team, 2021). Taxonomy assignment classification was performed with MiDAS5 database with increased coverage for bacteria and archaea (Dueholm et al., 2024). We used the method *assignSpecies* for taxonomic assignments at or near the resolution limit of the sequenced marker-gene that was developed specifically for the task of making species-level assignments to 16S data. ASVs that were classified as mitochondria, chloroplast, or eukaryota were removed. ASVs identified as contaminants were removed with *decontam* package (v2.20.0) (Davis et al., 2018) using extraction kit blank sequencing as negative controls. After quality filtering, six water samples were excluded from downstream analysis due to insufficient read counts.

*Metagenomic data analysis*. The sequenced reads were quality filtered using Trimmomatic (v0.40) to remove adaptors and low-quality sequences (Bolger et al., 2014). MetaPhlAn v4.1 (Blanco-Míguez et al., 2023) was used to assign taxonomy. Subsequently, the clean reads were *de novo* assembled into contigs via metaSPAdes (v3.15.5) (Nurk et al., 2017). Then, Maxbin (v2.2.7) was used to bin contigs (Wu et al., 2016) and checked for completeness and contamination using CheckM (Parks et al., 2015). These assemblies were annotated using Bakta (Schwengers et al., 2021), resistance gene identifier (RGI) v6.0.3 to determine ARGs with the CARD 4.0.0 database (Alcock et al., 2023), and geNomad v1.7.0 to annotate MGEs such as plasmids and viruses (Camargo et al., 2024). To calculate the abundance of annotated ARGs, FASTA sequences from RGI output were used for read pseudoalignment with Kallisto (v0.50.0). Then, the relative levels of ARGs from each sample were calculated separately as the mapped reads per kilobase per million reads (RPKM) (Bray et al., 2016). The MIMAG criteria (Bowers et al., 2017) was used to classify high quality (HQ)-MAGs as those with a completeness > 90% and contamination < 5%. HQ-MAGs were selected and used directly as they were generated using the workflow adopted in this study. Taxonomic annotation of MAGs was performed using the classify workflow from gtdb-tk v2.4.0 using the GTDB reference database (Chaumeil et al., 2020) and functionally annotated using Bakta v1.7. Co-localization of MGE and ARG was defined as the presence of a *qac* gene on a contig that also harbored either viral or plasmid-associated MGEs, based on GeNomad prediction.

### 2.6 Statistical analyses

All microbial community statistical analyses were conducted in R (v.4.3.3). For visualization of results, the packages ggplot2 (v3.5.1) and ampvis2 (v2.8.9) were used to generate heatmaps showing relative taxonomic abundance. Alpha diversity indices (Inverse Simpson) and richness estimators for the bacterial communities were calculated as described in the R package *phyloseq* (package version 1.44.0) software manual (McMurdie & Holmes, 2013). To test whether alpha diversity changed based on sink location, seasonality, or traffic, we calculated diversity indexes and performed significance testing by pairwise Wilcoxon tests for data not normally distributed. A p-value of <0.05 was considered statistically significant, with designations as follows: *\*\*\* p < 0.001; ** p < 0.01; * p < 0.05* and ‘ns’ non-significant. To generate ordination plots, we used principal coordinate analysis (PCoA) of Bray-Curtis distance to visualize variations in the microbial communities. To test for significant differences in the bacterial community structure, we performed permutational multivariate analyses of variance (PERMANOVA) using the ‘adonis’ function in the R package vegan v2.6-6.1 with 9999 random permutations within a given sink drain, across buildings and seasons. We also performed a similarity percentages breakdown (SIMPER) analysis to identify abundant ASVs that contributed most to the Bray-Curtis dissimilarity between buildings groups.

Spearman’s rank correlation was used to assess associations between alpha diversity and relative abundance, while Kendall’s tau correlation was applied in cases with a low number of samples to ensure robustness of the analysis.

### 2.7 Accession number and data availability

Raw amplicon sequence data are available in the SRA database under accession number PRJNA1254264 and BioSample accession numbers SAMN48105195 - SAMN48105473. Metagenomic reads were deposited under the same BioProject with BioSample accession numbers SAMN51280500 - SAMN51280527.

## 3. RESULTS AND DISCUSSION

We wanted to understand how spatial, seasonal, and traffic differences affect the microbial communities of sink drains. To do this, we collected and analyzed 288 biofilms (144 from each of the two buildings) and 18 composite water samples during high and low traffic across three seasons. An overview of the final sample set is shown in Table S1.

### 3.1. Four genera dominate the sink drain microbiome

The microbial communities in each academic building were characterized by two predominant genera, which together accounted for 30% of the composition. In the newer academic building, EH constructed in 2011, *Acinetobacter* (19.6 ± 10.4%) and *Enhydrobacter* (10.3 ± 10.1%) were the most abundant. In contrast, the microbial community in WC, constructed in 1960, was dominated by *Sphingomonas* (19.4 ± 19.7%) and *Mycobacterium* (10.4 ± 9.4%), with these genera being less abundant in EH, at 6.6% and 5.4%, respectively. Conversely, in WC, *Enhydrobacter* and *Acinetobacter* were present at lower relative abundances (6.9% and 4.5%, respectively) (Figure 1.a and b). The high relative abundance of *Mycobacterium* is similar to the findings from a study on sinks in dormitories in China (Hu et al., 2023), while *Sphingomonas* was also abundant in residential sinks assessed in Australia (Hayward et al., 2024) and showed increased abundance in women’s restrooms (Hu et al., 2023). Similarly, *Acinetobacter* was the most abundant across all sinks in a study performed on a university campus in the UK (Withey et al., 2021). Furthermore, *Acinetobacter* and *Enhydrobacter* were among the most abundant genera in the microbiome of residential kitchen sponges and surfaces across five European countries (Moen et al., 2023). It is important to note that *Enhydrobacter* and *Moraxella* are both members of the family *Moraxellaceae* and share 99.45% similarity in the 16S rRNA gene marker region, which may lead to potential mis-annotations between the two genera (Li et al., 2021). We therefore kept the assignment as *Enhydrobacter* but acknowledge the possibility that this taxon might be better assigned as *Moraxella*. Across all samples, other genera with a prevalence > 80% (> 230/288 samples) and a relative abundance > 1% included *Ottowia* (9.2%), *Microvirga* (8.9%), and *Brevundimonas* (3.2%) (Figure 1. a and b). These seven dominant and abundant genera together comprised the core microbiome of the sampled sink drains and seemed to align with similar environments around the world (Hayward et al., 2024; Hu et al., 2023; Moen et al., 2023; Withey et al., 2021; Withey & Gweon, 2024).

**Figure 1.**
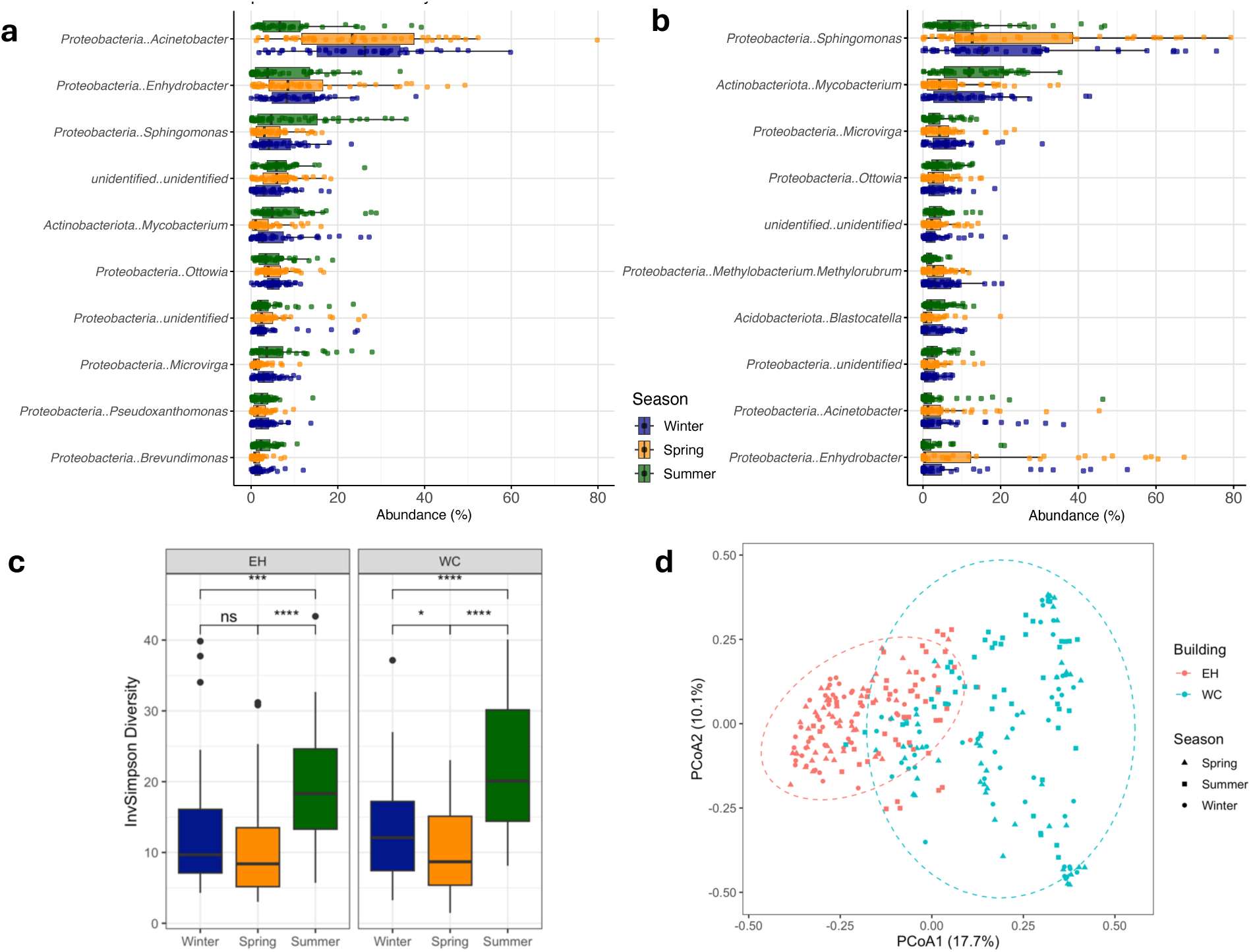
Microbial community composition of sink drain biofilms from academic buildings. Top panel displays boxplots of the top ten most abundant genera (with phylum-level classification added for reference) in the EH (a) and WC (b) buildings. Panel c visualizes the alpha diversity, invSimpson Diversity index across seasons for both EH and WC buildings. *\*\*\*\* p < 0.0001; *** p < 0.001; ** p < 0.01; * p < 0.05* and ‘ns’ non-significant. Panel d visualizes the PCoA plot showing the extent of variation in microbial communities with building and season as variable factors.

Contrary to the findings of Withey et al. (2021), where human skin flora was a dominant source of microbes for below-strainers sink communities, our study found the only predominant skin-associated taxa in the EH sinks were related to *Moraxella* (*Enhydrobacter*), a genus also linked to the human skin microbiome, as identified in the integrated Human Skin Microbial Gene Catalog (iHSMGC) (Li et al., 2021). In WC (an older building) our results showed that biofilms in high-water-usage areas like sinks are shaped more by environmental factors and were dominated by *Sphingomonas* and *Mycobacterium*. These genera are less directly related to human-associated microbiomes and more characteristic of disinfected water-adapted communities (Ma et al., 2020). The age of the WC building may reflect different building materials and a longer history of maintenance practices and disinfection protocols, potentially involving different products (e.g., chlorine or bleach-based products), which could have contributed to shaping its microbial community over time and selected for well-known chlorine disinfectant-resistant taxa such as *Mycobacterium* and *Sphingomonas* (Corinne et al., 2002; Galagoda et al., 2025; Sun et al., 2013). A previous study in a 50-years old building showed that architectural features, physical flow conditions, and chemical characteristics of premise plumbing influence the higher abundance the *Mycobacterium* in biofilms compare to bulk water ((Huang et al., 2021).

Sink biofilms exposed to BAC disinfectants daily had a community dominated by *Acinetobacter* (in the newer building, EH) and *Mycobacterium* (in the older building, WC). This suggests that a combination of building material and disinfectant exposure may enrich a specific set of taxa that persists in sink drain environments. Withey and Gweon (2024) demonstrated that a single sodium hypochlorite exposure led to a spike in *Acinetobacter* relative abundance from 3.16 to 74.26%. Although they observed a return to the pre-treatment community structure by the following week (Withey & Gweon, 2024), real-world systems encounter more frequent exposure, leading to long-term selection of certain taxa, as we observed here.

Despite the constant exposure to QAC disinfectants and surfactant soaps, the microbial diversity and composition of the biofilms revealed a complex prokaryotic community (Figure S1). Bacterial α-diversity varied across the sampled sink drains (Figure S2.A), but overall α-diversity indices were similar between the samples collected from the two buildings (Figure S2.B). The number of bacterial species co-inhabiting these sink drains was significantly higher in EH (136 ± 45), despite WC (112 ± 56) having been in service for 51 more years than EH (Figure S2.C). Seasonality significantly impacted bacterial diversity both between and within each building, with higher diversity in the summer compared to spring and winter (Figure 1.c). Unlike previous studies that reported a decrease in α-diversity over time (Mahnert et al., 2019; Withey & Gweon, 2024), our study spanned 230 days, from winter to the end of summer, and showed an increase of microbial diversity dependent on season (Figure 1.c).

### 3.2. Individual sinks, seasonality, and traffic explained the variability of microbial communities in academic buildings

We wanted to assess whether location, seasonality, and/or traffic influenced the microbial community structure. We observed noteworthy dissimilarities in the structure and composition of bacterial communities, with individual sink drains standing out as the most influential factor, explaining a substantial 43% (R^2^*_drain_* = 0.4309, F = 13.73, *p < 0.0001*) of microbial community variability (Figure 1.d). We also observed that the building and the sampling season significantly influenced the microbial community, although to a lesser degree (PERMANOVA, R^2^*_building_* = 0.1232, F = 40.12; R^2^*_season_* = 0.0313, F = 4.61; *p < 0.0001*). Moreover, when testing the interaction of the factors sink drain and season, the explained variability increased to 62% (PERMANOVA, R^2^*_drain*season_* = 0.6229, F = 8.43*; p < 0.0001*), suggesting that seasonal effects on microbial communities were highly sink-specific. Additionally, incorporating building occupancy (low vs. high traffic) within seasonal context for each sink had the highest explanatory power (R^2^*_drain*season*traffic_* = 0.7236, F = 5.29, *p < 0.0001*), indicating that the interaction among spatial, temporal, and anthropogenic factors plays a major role in shaping sink drain microbiomes.

Our findings align with and expand upon those of Withey et al. (2021), who analyzed drain biofilms on a university campus and found that each individual sink had the greatest impact on microbial community, while buildings (*n=9*) accounted for 19% of the variation.

We further determined whether differences existed between sinks A (the closest to the entrance) and B (the furthest from the entrance) of restrooms on the same floor across different levels of the buildings (Figure 2). The microbial communities varied, with 39% of the variation explained by sink location and floor level as factors in EH (PERMANOVA, R^2^*_drainAB*level_*= 0.386, F = 12.21, p < 0.0001) and 33% in WC (PERMANOVA, R^2^*_drainAB*level_* = 0.327, F= 9.42, p < 0.0001). The observed differences between sink A and B might be partly due to higher usage of sink A, likely influenced by the placement of paper towel dispensers near the entrance, which may have caused users to preferentially choose that sink.

**Figure 2.**
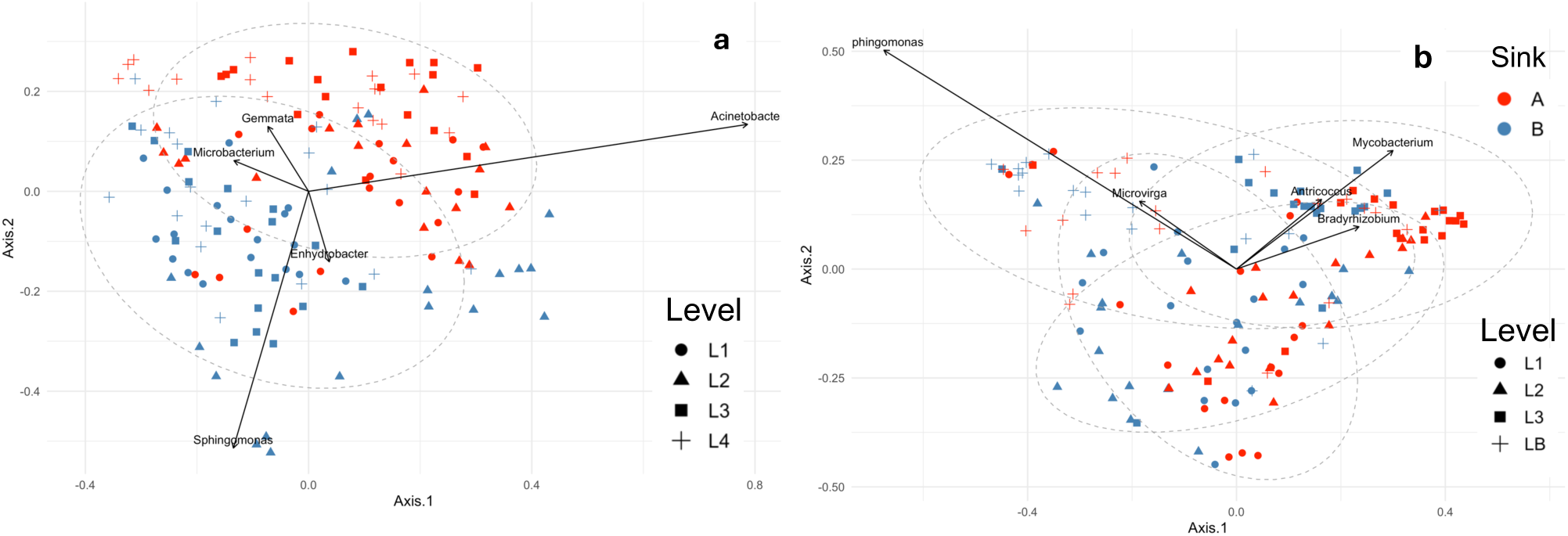
Microbial community of sink drain biofilms from academic buildings visualized as Bray-Curtis distances among samples in an ordination using multidimensional scaling for EH (a) and WC (b).

We identified five highly abundant ASVs that significantly contributed to the dissimilarity between sink groups. In EH, the presence of *Acinetobacter*, accounting for 12% of the dissimilarity, highly influenced the microbial communities of the biofilm found in A sinks (Figure 2.a). On the other hand, *Enhydrobacter* and *Sphingomonas* were the most discriminant species associated with B sinks, located furthest from the entrance (Figure 2.a).

In the WC academic building, where the floor distribution of restrooms explained 22% of the variation in the structure of the microbial communities, *Sphingomonas sp*. and *Microvirga* were the ASVs contributing most to the dissimilarity observed in the basement (LB), whereas *Mycobacterium* and *Bradyrhizobium* were associated with the third floor (L3) restroom (Figure 2.b). The genus *Sphingomonas* was strongly associated with the basement floor in WC, which had the lowest occupancy and consequently the least frequent use, which was also the case with the B sinks in EH. These associations suggest a link with low-traffic conditions and an increased relative abundance during stagnation periods, as observed in a previous study (Calero Preciado et al., 2021).

### 3.3. Microbial community of the tap water differed from those in sink-drain biofilms

We collected bulk composite water samples during each sampling campaign to assess whether microbial growth in the sink is shaped by or seeded from the waterborne microbial community. The composite water samples exhibited a complex and diverse bacterial community, consisting of up to 835 ASVs distributed across 18 phyla. The two most prevalent phyla were *Proteobacteria* (currently known as *Pseudomonadota*) (55.7%) and *Cyanobacteria* (10.9%), with similar abundances in both buildings. Within the *Proteobacteria* phylum, the two most abundant classes were *Alphaproteobacteria* and *Gammaproteobacteria*. In contrast to the biofilm phase (Fig S1), *Alphaproteobacteria* were more predominant in the EH building (39.5%) compared to WC (22.6%), whereas *Gammaproteobacteria* were more prevalent in WC (29.8%) than in EH (19.5%). This profile indicates a microbial community composition in the water phase that contrasts with that observed in the sink biofilm phase (Figure 3.a). Importantly, genera that were highly abundant in the sink biofilm phase, such as *Sphingomonas, Acinetobacter, Enhydrobacter, Mycobacterium*, were present at less than 6.5% in the bulk water phase (Figure 3.a).

**Figure 3.**
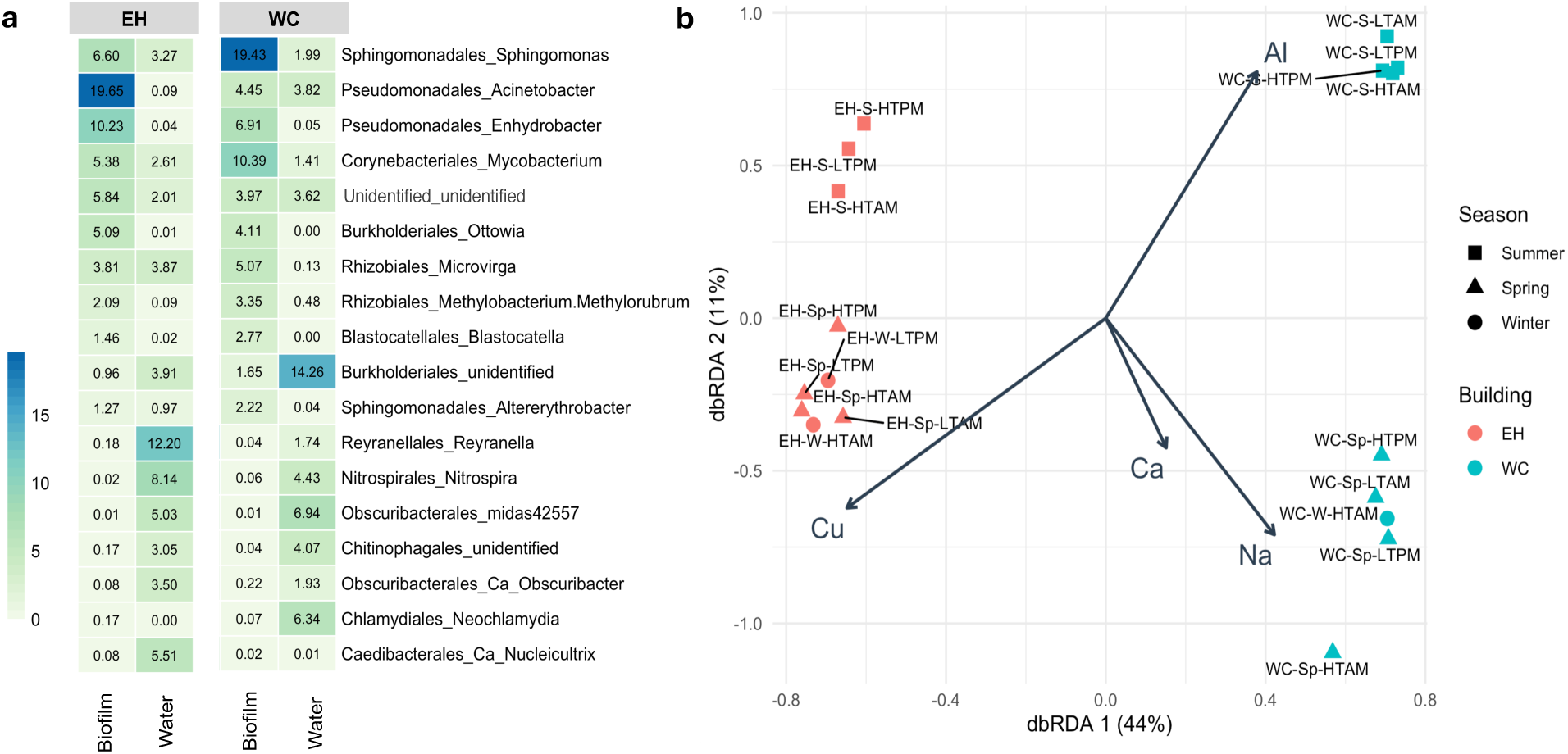
Heatmap depicting the percent relative abundance of microbial communities at the genus level across water and biofilm phases in each sampled building (EH and WC). The mean relative abundance of the top 10 most dominant genera from each phase were selected, with taxonomic order label added for clearer reference (a). Distance-based redundancy analysis (dbRDA) of microbial community composition at the ASV level (using Hellinger transformation) from sink water samples and water chemical quality (b). The vectors represent selected chemical factors: Cu, copper; Ca, calcium; Na, sodium; Al, aluminum. Cobalt, vanadium, molybdenum, iron, manganese, silver, zinc, and nickel were excluded due to high collinearity (Table S3).

The differences in microbial communities between EH and WC buildings could be partially explained by variations in water chemistry, likely due to different building materials. Samples from EH exhibited higher concentrations of copper, cobalt, nickel, and zinc, particularly during spring and winter compared to summer (Figure 3.b, Table S3). EH is a newer building (14 years in service), and its premise plumbing system may be leaching metals from the copper pipes into the water phase, acting both as an antimicrobial agent and as a micronutrient. Consequently, it may influence the abundance of *Acinetobacter* and other bacteria sensitive to copper (Williams et al., 2020) while also promoting the growth of genera *Reyranella* and *Nitrospira* (van der Kooij et al., 2020). In contrast, water from WC had higher concentrations of sodium, calcium, and magnesium during colder months, and elevated levels of aluminium, beryllium, and arsenic during the warmer season (Figure 3.b). Aluminium concentrations may vary seasonally because the solubility of aluminium hydroxide, commonly used during coagulation treatment, is temperature dependent above pH 6-7, typical of drinking water (Trueman et al., 2022).

None of the samples analyzed exceeded permissible levels of metals according to drinking water quality standards (EPA, 2025).

In the water phase of the EH building, microbial diversity increased with high traffic (Figure S3.A), similar to observations by Greenwald et al. (2022), who found that frequent water flow and reduced stagnation promote microbial diversity in building water systems. During high-traffic periods, water usage likely introduces a broad range of microbial taxa into the water phase, enriching the community through a constant influx of microorganisms from building plumbing and environmental exposure. In contrast, during low-traffic periods when water stagnation is more common, diversity in the water phase decreases (Figure S3.A), likely due to limited microbial inputs and the dominance of a few resilient taxa adapted to stagnant conditions. Interestingly, in the biofilm of the sinks located closer to the door (A), presumably with higher usage, we observed the opposite trend: microbial diversity was higher during low-traffic periods and decreased when traffic was high (Figure S3.B). This inverse relationship could be due to the selective pressures imposed by frequent handwashing and soap use during high-occupancy times. Soaps and disinfectants tend to reduce biofilm diversity by selecting for a few resistant species, such as *Acinetobacter* and *Mycobacterium*, which possess efflux pump systems and outer membrane structures that may enable them to survive under these conditions (Singh et al., 2017).

In the case of some Gram-negative bacteria, the lipopolysaccharide layer of the outer membrane acts as a barrier, offering intrinsic resistance to QACs by preventing interaction with the cytoplasmic membrane (Maillard & Pascoe, 2024). In addition, the genus *Mycobacterium* (phylum *Actinobacteria*), one of the predominant taxa in all of our samples, possesses a lipid-rich outer layer of mycolic acids surrounding the cell that makes it one of the least susceptible to biocides (Lambert, 2002). In contrast, many of the genera that are more abundant in the water phase, such as *Reyranella* and *Nitrospira*, may be more susceptible to QAC disinfectants and unable to persist in the biofilm phase due to these selective pressures (Zhang et al., 2023). When exposed to BAC and surfactants, these genera might experience cell membrane disruption and cytoplasmic leakage, preventing their establishment in biofilms (Mc Carlie et al., 2020; Tong et al., 2021). The resistant taxa could outcompete the more susceptible organisms within the biofilm, leading to a decline in overall community diversity. Conversely, during low-traffic periods, fewer direct inputs from human contact and fewer disinfection events may allow a wider array of microbial species to coexist within the biofilm, supporting higher diversity. These findings suggest that biofilm microbial communities respond to inputs from human activity, disinfection practices, and water dynamics differently than the free-floating communities in the water phase.

### 3.4. Acinetobacter linked to usage pattern and inversely correlated with microbial diversity

Among the 18 ASVs associated with the genus *Acinetobacter,* ASV2 was the most prevalent and abundant. ASV2, which was the only one assigned as *Acinetobacter parvus,* appears to be related to usage patterns in EH. *Acinetobacter* was significantly more abundant during periods of high traffic, when students were in session, compared to the academic break periods in spring and summer (Figure S4.A). Moreover, the A sinks displayed distinct patterns in response to varying levels of traffic. There was a significant correlation between the relative abundance of *Acinetobacter* in sinks A and the occupancy count of the EH building during the summer season (Figure 4.a). The first floor of WC, the most frequently used area with higher traffic, also showed a higher relative abundance of *Acinetobacter* compared to less frequently trafficked floors (Figure S4.B). Overall, low traffic (fewer than 20 users per restroom), had a negative impact on the relative abundance of *Acinetobacter* (Figure S5). The observed higher relative abundance of *Acinetobacter* during periods of increased traffic underscores the role of anthropogenic disturbance in shaping microbial community composition within built water-associated environments. Rather than eliminating microbial communities, frequent usage and cleaning may create selective conditions favouring stress-tolerant taxa enrichment with the capacity to persist under disinfectant exposures (Tandukar et al., 2013).

**Figure 4.**
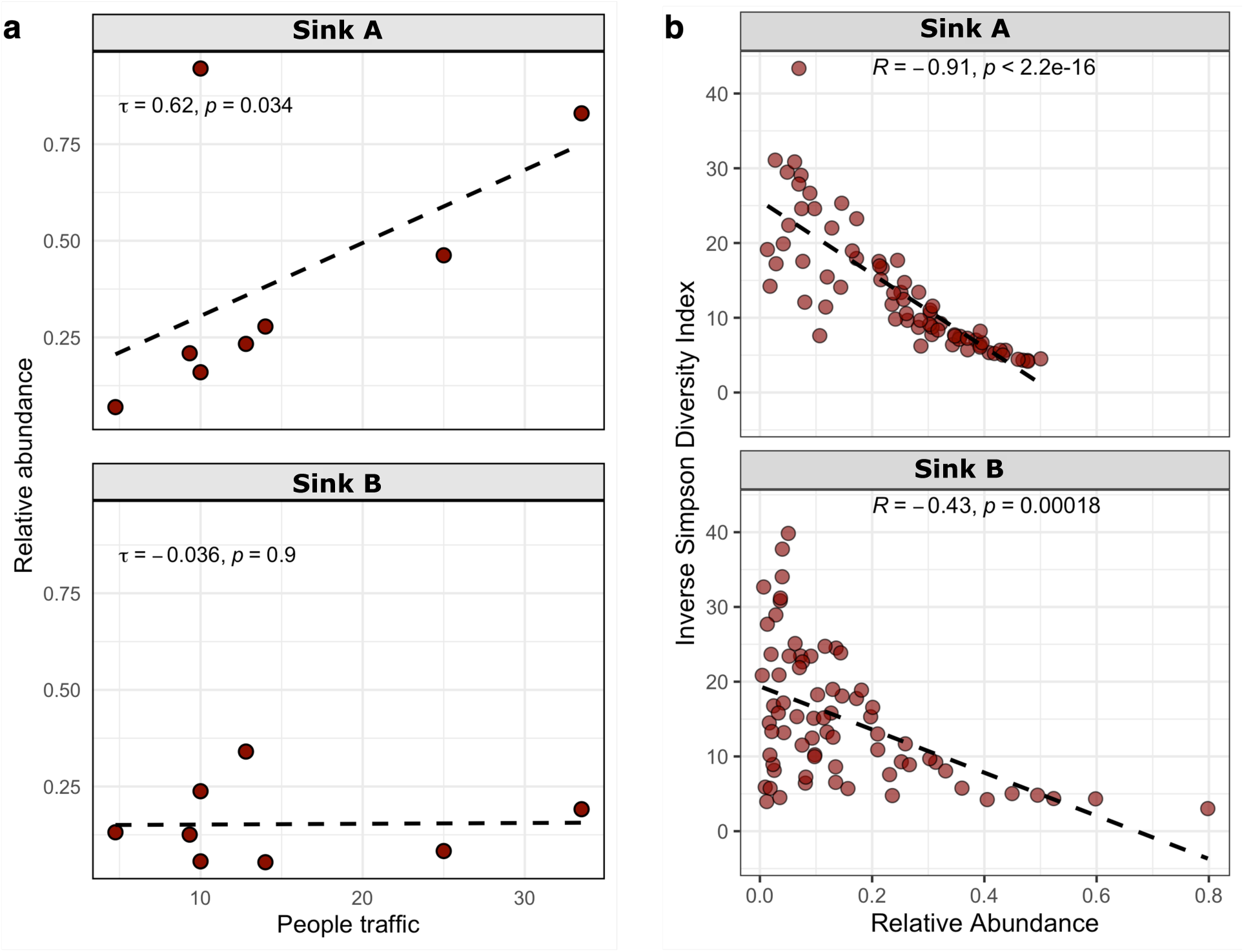
a. Kendall correlation between the relative abundance of the genus *Acinetobacter* in A sinks (closest to the restroom entrance) and B sinks (furthest to the entrance) of EH and traffic during the summer season (a). Spearman correlation between genus *Acinetobacter* relative abundance and Inverse Simpson α-diversity index of A and B sinks in EH (b).

Furthermore, *Acinetobacter* showed a strong inverse correlation with microbial diversity, indicating that it often dominates the sink drain niche (Figure 4.b). High relative abundance and prevalence of *Acinetobacter* in sinks have been previously reported (Withey et al., 2021).

Moreover, a recent study documented a peak in *Acinetobacter* abundance following a single disinfection with sodium hypochlorite (Withey & Gweon, 2024). These outcomes suggest that the genus *Acinetobacter* is enriched in frequently disinfected environments.

These findings contribute to the increasing body of evidence that regular exposure to biocidal treatments may not achieve the desired level of disinfection (Boyce, 2023; J. Maillard, 2022; Mc Carlie et al., 2020; Merchel Piovesan Pereira & Tagkopoulos, 2019). Instead, frequent cleaning regimes and human activity may shape the microbial communities, favoring microbes that are resistant to the employed disinfection protocols, ultimately leading to more resilient microbial communities (Tandukar et al., 2013). Although these communities are located in sink drains, rather than on frequently touched surfaces, their relevance lies in their potential to serve as reservoirs of microbes, which can spread and propagate from drains via aerosolization during faucet use, thereby contributing to the overall microbial load within indoor spaces (Döring et al., 1996). In healthcare settings, it has been reported that during sink use, droplets carrying carbapenem-resistant *Pseudomonas aeruginosa* can be released from sink drains, thus contaminating the surrounding environment (Cahill et al., 2023). Furthermore, a high percentage of isolates from sinks/sink drains have been found to be genetically linked to clinical strains, supporting evidence of transmission from environmental reservoirs to patients (Choquet & Mullié, 2022).

### 3.5 Evidence for disinfectant resistance genotypes and corresponding phenotypes

Because the sinks were disinfected with QACs daily, we wanted to understand the impact this would have on the antibiotic resistance gene profile of the sink microbiomes. Accordingly, we sequenced shotgun metagenomes from four sink drain biofilm samples - two from each building with contrasting occupancy. Samples were collected during classes in late summer (second week of the fall semester), representing high traffic on the first floor (EH and WC), and low traffic on the third floor (EH) and basement (WC) (Table S2).

Seven high-quality (> 95% completeness and < 5% contamination) metagenome-assembled genomes (MAGs) were recovered from the metagenomic sequence reads, adhering to the Minimum Information about a Metagenome-assembled Genome (MIMAG) standards (Bowers et al., 2017) for bacteria and archaea (Supplemental Table S4). The most abundant genera identified by 16S rRNA amplicon sequencing were also represented among the recovered MAGs. *Acinetobacter parvus*, prominent in the 16S dataset, was recovered as a MAG from a WC sample. Similarly, *Mycobacterium*, one of the dominant genera in both buildings, was represented by two high-quality MAGs: *M. phocaicum* and *Mycobacterium* sp., both non-tuberculous species outside the *Mycobacterium avium*-intracellular complex (MAC), recovered in EH and WC samples, respectively. Although the genus *Nakamurella* was not detected in our 16S dataset, we recovered a high-quality MAG closely related to *Nakamurella flava*. This taxon was of note because it was previously reported as a predominant organism in dormitory drain pipes in China (Hu et al., 2023). Medium-quality MAGs (76.84 % completeness and 4.25% contamination) assigned as *Moraxella sp902506215* were recovered from samples in both buildings. It is important to note that *Moraxella sp902506215,* classified using the GTDB-tk database for MAGs taxonomy assignment (Chaumeil et al., 2020), corresponds to *Enhydrobacter* UBA1882 in the NCBI taxonomy, confirming that *Moraxella* is one of the predominant genera in our sink microbiome samples.

Across all samples, a total of 29 distinct ARGs were identified (Figure S6), 14 of which were found among the seven HQ-MAGs (Figure 5). Of the 29, we found four genes responsible for QAC resistance (Figure S6), specifically those in the *qac* operon (Cervinkova et al., 2012; Smith et al., 2008). *A. parvus*, *Ottowia sp*. and *Amaricoccus sp*. were the only three MAGs identified to carry genes implicated in conferring resistance to QAC-based disinfectants (Figure 5). *A. parvus* carried two *qacG* genes, one of which was co-localized in a contig carrying plasmid sequences, and a *rsmA* gene, which is linked to adhesion in biofilm formation, desiccation resistance, and potentially enhanced multidrug resistance (Farrow III et al., 2020; Zhao et al., 2023). *Ottowia sp.* MAG had three *qac* genes (*qacL, qaG* and s*ugE*) and two *adeF* genes, subunits of the intrinsic resistance- nodulation-(RND) cell division efflux pump AdeFGH, which can confer resistance to fluoroquinolone and tetracycline (Sébastien et al., 2010). Among all samples, *qacG* and *adeF* were the two most abundant ARGs with highest abundance observed in samples from the first floors of both buildings (Figure S6). This pattern may be attributed to higher traffic on the first floors, resulting in greater microbial input into the sink biofilms. Such findings are consistent with prior research on microbial diversity in high-use environments, where ARG prevalence is influenced by occupancy and sanitation practices (Mahnert et al., 2019). Vancomycin resistance genes (*vanY, vanW, and vanT*) were also frequently detected in all samples (Figure S6) and were identified in four HQ-MAGs, including those classified as *Mycobacterium* and *Chryseobacterium* recovered from the WC building. Exposure to metals, particularly Al and As, has been associated with vancomycin resistance in other organisms (Sadeghian et al., 2024), and could be why *van* genes were found in the WC building (Table S3). Two *Mycobacterium* MAGs carried the *murA* gene, which confers resistance to fosfomycin.

**Figure 5.**
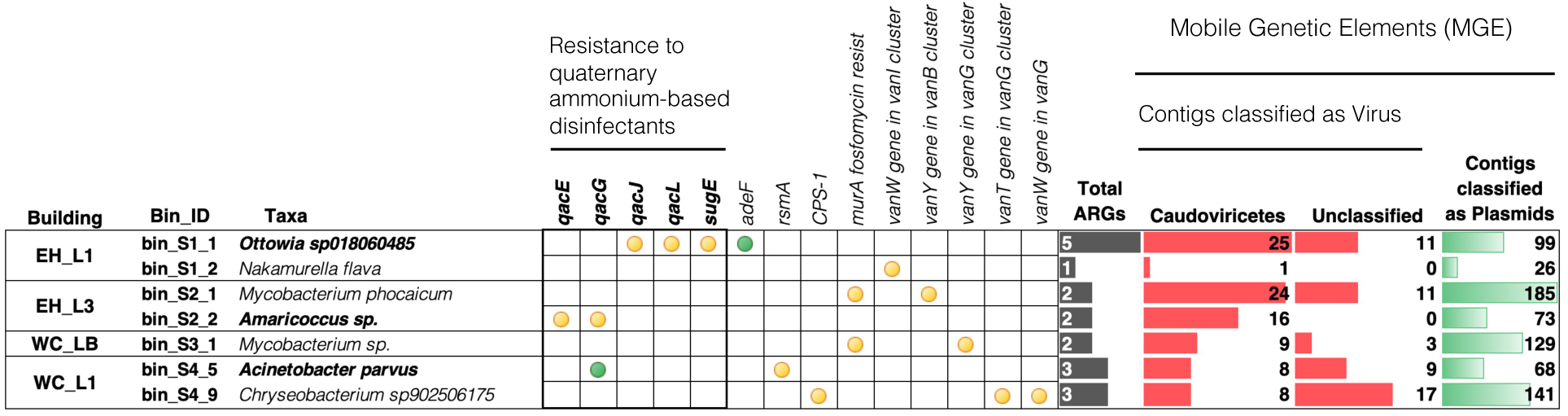
Antibiotic and disinfectant resistance genes (perfect and strict matching criteria) and mobile genetic elements (virus and plasmids) annotated on HQ-MAGs. High traffic (L1) and low traffic (L3 and LB) of two buildings (EH and WC) are shown. The green circles indicate the presence of two genes, while yellow circles indicate the presence of one gene.

We next wanted to link the potential QAC-resistant genotypes (Figure 5) with phenotypes associated with reduced susceptibility to QACs. We assessed the phenotypes by plating microbes from sink drain swabs on increasing concentrations of BAC-containing agar media (Figure S7 and S8). Concentrations of BAC above 100 mg/L reduced the culturable microbial community to 20% of the total (Figure S9). We observed growth of *A. parvus* at all concentrations of disinfectants tested (1 – 100 mg/L BAC and commercial Virex, Figure S7 and S8). The methodology of determining minimum inhibitory concentration (MIC) can alter the actual value, and therefore, make cross-method comparison challenging; however, previous studies have reported the MIC of different clinical *Acinetobacter* isolates to be between 50 and 100 mg/L BAC (Kawamura-Sato et al., 2008).

MAGs assigned as *Ottowia* and *Moraxella* were recovered from plates with 100 mg/L and low concentrations of Virex (Table S5.1). Notably, the *Ottowia-*MAG contained a contig with *qacL* genes. Several species of the genus *Chryseobacterium* (sp902506175, *C. formosense, C. aquaticum, C. mucoviscidosis*) were recovered from plates containing high concentrations of disinfectants (100 mg/L of BAC and Virex) (Table S5.1). It is worth noting that a previous study reported that disinfectants containing BAC could be more effective than chloroxylenol or chlorhexidine gluconate disinfectants at removing *Chryseobacterium* species from contaminated surfaces (Mwanza et al., 2022). Furthermore, the presence of the *vanT* gene in *vanG* cluster (Figure 5) links vancomycin resistance to biofilm formation, which may contribute to reduced susceptibility to disinfectants (Patra et al., 2025).

In addition, MAGs belonging to the genera *Sphingomonas (S. adhaesiva and S. hankookensis_A)* and *Brevundimonas (B. colombiensis, B. diminuta_C, B. huaxiensis, B. pondensis),* which were among the most abundant taxa in the 16S dataset, were also recovered from plates containing high disinfectant concentrations (Figure S7 and S8, Table S5.2). *Brevundimonas spp*., in particular *B. diminuta*, is a genus of non-fermenting Gram-negative bacteria considered as an emerging global opportunistic pathogen associated with clinical bacteraemia (Ryan & and Pembroke, 2018).

The MAGs classified as *Acinetobacter, Moraxella,* and *Chryseobacterium* that were abundant in the 16S dataset and cultured on BAC-containing agar belong to a group known as non-fermentative Gram-negative rods (Vaneechoutte et al., 2015). These genera include species related to risk group 2, which poses moderate individual risk but low community risk (CDC, 2019). *A. parvus* is frequently isolated from blood cultures, with infections related to intravascular catheters (Nemec et al., 2003). *Moraxella* species are rare agents of infections (conjunctivitis, keratitis, otitis) and are also known to colonize the upper respiratory tract (Vaneechoute et al., 2000). Furthermore, the species *M. osloensis* is recognized as a cause of bad laundry odor (Kubota et al., 2012). The genus *Chryseobacterium* is an uncommon human pathogen (Tez and Tez, 2018), but some species possess inherent resistance to multiple antibiotics, including carbapenems and cephalosporins (Parajuli et al., 2023), and have emerged as opportunistic pathogens causing hospital-acquired infections (Poonam et al., 2015). Notably, all three genera have been identified as frequent colonizers of kitchen sponges and surfaces across different countries (Cardinale et al., 2017).

### 3.6 Potential horizontal gene transfer (HGT) and ARG dissemination

Our findings revealed a notable prevalence of QAC resistance genes (*qacG, qacL, qacJ, qacE*), with *qacG* being the most frequently detected across all samples and co-localized with contigs carrying plasmid genes (plasmid-contig length range 493-96,633 bp, 60% of contigs < 5000 bp, Figure S10). Between 34% and 53% of *qac* genes were co-localized with contigs associated with mobile plasmids and viruses, with the sample from the first floor of WC showing the highest percentage of possible plasmid-carrying *qac* genes. The *qac* genes, encoding resistance to QACs, are known to be plasmid-borne genes (Bjorland et al., 2003). Biofilms provide a niche for more resistant bacterial communities, as biofilm-dwelling bacteria withstand higher concentrations of disinfectants due to reduced penetration and prolonged exposure. Previous studies have shown that if a disinfectant is not properly prepared or used at a subinhibitory concentration, it may select for a resistant population, potentially driving resistance to both disinfectants and antibiotics (Lineback et al., 2018; Maillard & Centeleghe, 2023; Nordholt et al., 2021; Tong et al., 2021).

This supports previous research suggesting that regular use of QAC-based disinfectants can exert selective pressure, enabling QAC-resistant bacteria to persist in these environments (Maillard & Pascoe, 2024; Minjae et al., 2018). The co-location of *qac* genes on plasmids and viral elements further underscores the potential for HGT, facilitating ARG dissemination within and beyond the sink drain biofilms. The presence of MGEs, such as plasmids associated with *qac* genes, suggests an increased risk of ARG spread in high-use public buildings, a finding consistent with recent studies that highlight the role of MGEs in ARG mobility (Mahnert et al., 2019).

## CONCLUSIONS

Our study revealed that location and seasonal variations significantly shaped microbial community composition in sink drains. Additionally, we found that sink usage patterns mediated the prevalence of ARGs, including *qac* genes, as well as the relative abundance of certain bacterial taxa, particularly *Acinetobacter* species.

*Acinetobacter* relative abundance positively correlated with increased traffic and exhibited an inverse relationship with microbial diversity, suggesting environmental factors and human activity shaped the microbiome. The persistence and abundance of *Gammaproteobacteria* such as *Acinetobacter, Moraxella (Enhydrobacter),* and *Chryseobacterium,* is consistent with previous findings in highly disinfected environments (Cardinale et al., 2017; Moen et al., 2023), similar to those in hospitals and residential kitchens.

The high prevalence of *qac* genes on plasmids, particularly in *Acinetobacter,* highlights the potential for resistance gene dissemination through HGT, emphasizing the need for targeted strategies to mitigate risks in built environments. Controlling bacteria in sink drains presents a significant challenge due to their ability to form biofilms that are highly resistant to disinfectants. The frequent use of disinfectants, especially QACs, in these settings may exert selective pressure, favoring the survival of bacteria that harbor ARGs.

Our findings offer an understanding of the sink bacterial communities and might serve as a baseline for future surveillance efforts monitoring community stability and resilience in the built environment. Further research on the metabolic functions of microbes in the built environment and factors influencing communal sink drain biofilms will be essential to improve our ability to manage and control microbial communities in highly disinfected settings, reduce potential health risks, and ultimately foster healthier indoor environments.

## ACKNOWLEDGMENTS

This work was supported by the U.S. Department of Defense, Office of Army Research under Grant number W9132T-22–2-0001 and W9132T-23–2-0003. We thank Mike Dollhopf for ICP-MS analyses. Any opinions, findings, and conclusions or recommendations expressed are those of the authors and do not necessarily reflect the official policies or positions of the U.S. Army Corps of Engineers/ERDC, the Department of Defense, or the U.S. Government.

K.R.H., M.C.C, B.M., and C.W.M. conceptualized the study. M.C.C., J.L., S.B., R.R., N.M.B., R.M, and B.R. performed sampling, conducted the investigation and laboratory analysis. M.C.C. performed the data analysis and its visualization and wrote the original draft. C.W.M. B.M., and K.R.H. helped interpreting the results, supervision, and funding acquisition. All authors edited and approved the final manuscript.

